# Strand-biased circularizing integrative elements spread *tmexCD-toprJ* gene clusters encoding RND-type multidrug efflux pumps by repeated transpositions

**DOI:** 10.1101/2022.09.22.508988

**Authors:** Trung Duc Dao, Hirokazu Yano, Taichiro Takemura, Aki Hirabayashi, Le Thi Trang, Hoang Huy Tran, Keigo Shibayama, Futoshi Hasebe, Ikuro Kasuga, Masato Suzuki

**Affiliations:** Vietnam Research Station, Center for Infectious Disease Research in Asia and Africa, Institute of Tropical Medicine, Nagasaki University, Nagasaki, Japan; Antimicrobial Resistance Research Center, National Institute of Infectious Diseases, Tokyo, Japan; National Institute of Hygiene and Epidemiology, Hanoi, Vietnam; Nagoya University Graduate School of Medicine, Nagoya, Japan; Vietnam-Japan University, Vietnam National University, Hanoi, Vietnam; Research Center for Advanced Science and Technology, The University of Tokyo, Tokyo, Japan

**Author notes:** Contributed equally (Trung Duc Dao, Hirokazu Yano). Corresponding authors. E-mail addresses (I. Kasuga); (M. Suzuki).

**Keywords:** strand-biased circularizing integrative element, SE, *tmexCD-toprJ*, tigecycline, *Aeromonas*

## Abstract

Antimicrobial resistance genes (ARGs) are associated with mobile genetic elements (MGEs) that conscript useful genes into the human–microbe and microbe–microbe battlefields. Thus, under intense selective pressure, ARGs have been constantly adapting and evolving, spreading among microbes. *tmexCD-toprJ* gene clusters, which encode resistance–nodulation–cell division (RND)-type efflux pumps, confer multidrug-resistance to clinically important antimicrobials, including tigecycline. Noteworthily, these gene clusters have emerged in gram-negative bacteria in humans, animals, and the environment worldwide by MGE-mediated transfer. Here we show a hidden MGE, strand-biased circularizing integrative element (SE), that is recently recognized to mediate transpositions of ARGs, associated with the spread of *tmexCD-toprJ* gene clusters. We identified multidrug-resistant isolates of *Aeromonas* species in a water environment in Vietnam that harbored multiple copies of *tmexCD-toprJ* in their chromosomes that were associated with SEs. In particular, *Aeromonas hydrophila* NUITM-VA1 was found to harbor two copies of a novel variant of *tmexC3*.*3D3*.*3-topJ1* within cognate SEs, whereas *Aeromonas caviae* NUITM-VA2 harbored four copies of a novel variant of *tmexC2D2*.*3-topJ2* within cognate SEs. Based on the nature of SE to incorporate a neighboring sequence into the circular form and reinsert it into target sites during transposition, we identified the order of intragenomic movements of *tmexCD-toprJ* gene clusters. Altogether, our findings suggest that most known subgroups of *tmexCD-toprJ* and their subvariants underwent transpositions among bacterial chromosomes and plasmids via SEs. Hence, a *tmexCD-toprJ* gene cluster ancestor may have been initially mobilized via SE, subsequently spreading among bacteria and evolving in new hosts.

## Introduction

Antimicrobial resistance genes (ARGs) are estimated to have originated in environmental bacteria and subsequently spread to pathogenic bacteria in humans as acquired ARGs (1). Mobile genetic elements (MGEs) are the major driving force of gene transfer and amplification underlying ARG evolution (2). Spontaneous mutations and antimicrobial selection have led to the emergence of clinically problematic ARGs, including carbapenemase genes (*e*.*g. bla*_NDM_, *bla*_KPC_, *bla*_OXA-48-like_, *bla*_IMP_, *bla*_VIM_, and *bla*_GES-5-like_) (3), colistin resistance genes (*e*.*g. mcr* and *arnT*) (4, 5), and tigecycline resistance genes [*e*.*g. tet*(X) and *tmexCD-toprJ*] (6).

Carbapenem, colistin, and tigecycline are considered last-resort antimicrobials for infections caused by multidrug-resistant (MDR) gram-negative bacteria, such as *Enterobacterales, Pseudomonas aeruginosa*, and *Acinetobacter baumannii*, which are serious global public health threats (3, 4, 6). *tet*(X4) and other *tet*(X) variants, which encode flavin-dependent monooxygenases that catalyze tigecycline degradation, have emerged mainly in *Enterobacterales* and *Acinetobacter* species (3). *tmexCD1-toprJ1, tmexCD2-toprJ2, tmexCD3-toprJ1* (initially designated *tmexCD3-toprJ3*), *tmexCD4-toprJ4*, and other *tmexCD-toprJ* subvariants, which encode resistance–nodulation–cell division (RND)-type efflux pumps that excrete multiple antimicrobials, including tigecycline, have emerged worldwide mainly in *Enterobacterales* and *Pseudomonas* species (3, 7–16).

Recently, a novel MGE family, named strand-biased circularizing integrative element (SE), was identified in *Vibrio alfacsensis* (17) and later in more diverse taxa of *Gammaproteobacteria* (18). SEs move between genomic locations via a copy-out-like route using tyrosine recombinases: SEs incorporate 6 bases next to the 5′-end of one specific strand into their circular intermediate form, which are then reinserted into a new target location. Therefore, SE-mediated transposition results in the insertion of 6 bases at newly formed *attR* recombination sites (17), and this 6-bp footprints allow tracing the order of SE integration events that occurred in the past in a genome.

In this study, we investigated MDR bacteria in a water environment in Vietnam and found MDR isolates of *Aeromonas* species that harbored multiple genes conferring resistance to last-resort antimicrobials, including *tmexCD-toprJ* gene clusters. Close examination of MGEs associated with multiple copies of *tmexCD-toprJ* in the chromosomes indicated involvement of SEs in the intragenomic amplification of the gene clusters. Comparison of sequences around known variants of *tmexCD-toprJ* in publicly available genome data revealed that most of them were likely integrated into bacterial chromosomes and plasmids via SEs.

## Results and Discussion

Two carbapenem- and tigecycline-resistant isolates of *Aeromonas* species, namely NUITM-VA1 and NUITM-VA2, respectively, were obtained from a water environment in Vietnam in 2021. Whole-genome sequencing analysis revealed that NUITM-VA1 (accession no. <u>AP025277</u>) and NUITM-VA2 (accession no. <u>AP025280</u>) were 97.1% and 97.9% identical to *Aeromonas hydrophila* strain ATCC 7966^T^ (accession no. <u>CP000462</u>) and *Aeromonas caviae* strain CECT 838^T^ (accession no. <u>JAGDEN000000000</u>), respectively. Multilocus sequence typing analysis revealed that NUITM-VA1 and NUITM-VA2 belonged to novel sequence types (STs), ST1063 and ST1064, respectively, of *Aeromonas* species.

*A. hydrophila* NUITM-VA1 harbored multiple clinically important ARGs, such as intrinsic genes of cephalosporinase (*bla*_cepH_) and oxacillinase (*bla*_aphH_), tigecycline resistance genes [*tet*(X4) and *tmexCD3-toprJ1*-like], phosphoethanolamine transferase gene conferring colistin resistance (*mcr-3*.*9*), and efflux pump gene conferring fluoroquinolone resistance (*qnrVC4*), whereas *A. caviae* NUITM-VA2 harbored carbapenemase genes (*bla*_NDM-1_, *bla*_KPC-2_, and *bla*_VIM-4_), tigecycline resistance genes (*tmexCD2-toprJ2*-like), and 16S ribosomal RNA methyltransferase gene conferring aminoglycoside resistance (*rmtB*) (Figs. 1A and B). Noteworthily, NUITM-VA1 and NUITM-VA2 harbored multiple copies of tigecycline resistance genes. Consistent with this observation, NUITM-VA1 and NUITM-VA2 showed low susceptibility to most antimicrobials tested, including carbapenems, cephalosporins, aminoglycosides, fluoroquinolone, and colistin with some exceptions (Table 1).

**Fig. 1.**
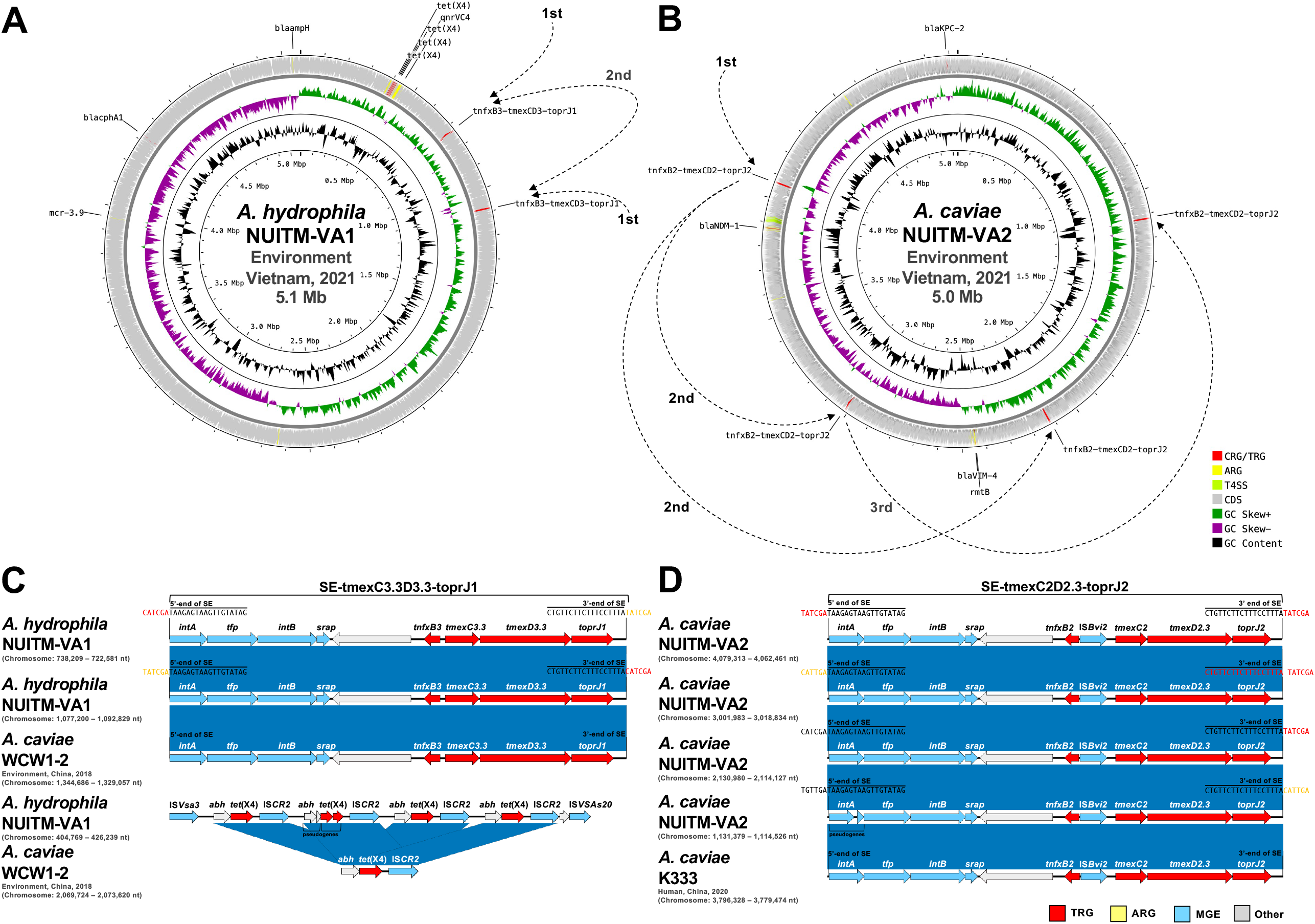
(A and B) Circular representation of chromosomes of *A. hydrophila* NUITM-VA1 (accession no. <u>AP025277</u>) (A) and *A. caviae* NUITM-VA2 (accession no. <u>AP025280</u>) (B) isolated in Vietnam in 2021 in this study. The dashed arrows indicate the putative order of SE-mediated transpositions of *tnfxB-tmexCD-toprJ*. (C and D) Linear comparison of *tmexCD-toprJ*-containing SEs and *tet*(X4)-containing regions in *A. hydrophila* NUITM-VA1 (C) and *A. caviae* NUITM-VA2 (D) with those of *A. caviae* WCW1-2 (accession no. <u>CP039832</u>) isolated from an environment in China in 2018 (19) and *A*.*caviae* K333 (accession no. <u>CP084031</u>) isolated from a human in China in 2020 (20). 5’- and 3’-ends of SEs and their 6-bp transposition footprint sequences in *Aeromonas* species isolates in this study are shown. Red, yellow, light green, light blue, gray, green, purple, and black indicate carbapenem and tigecycline resistance genes (CRG and TRG), other antimicrobial resistance genes (ARG), mobile gene elements (MGE), type IV secretion system-associated genes involved in conjugation (T4SS), other coding sequences (Other), GC Skew+, GC Skew–, and GC content, respectively. The blue color in comparison of sequences indicates almost 100% identity.

**Table 1.** MICs of antimicrobials against *A. hydrophila* NUITM-VA1 and *A. caviae* NUITM-VA2. The efflux pump inhibitor 1-(1-naphthylmethyl)-piperazine (NMP) was used at 75 mg/L. Abbreviations: TIG, tigecycline; MIN, minocycline; DOX, doxycycline; TET, tetracycline; IPM, imipenem; MEM, meropenem; CTX, cefotaxime; CAZ, ceftazidime; CIP, ciprofloxacin; AMK, amikacin; GEN, gentamicin; TOB, tobramycin; STR, streptomycin; CST, colistin.

| Isolate | MIC (mg/L) |  |  |  |  |  |  |  |  |  |  |  |  |  |
| --- | --- | --- | --- | --- | --- | --- | --- | --- | --- | --- | --- | --- | --- | --- |
|  | TIG<br>(+NMP) | MIN | DOX | TET | IPM | MEM | CTX | CAZ | CIP | AMK | GEN | TOB | STR | CST |
| *A. hydrophila*<br>NUITM-VA1 | 64<br>(8) | 64 | 128 | >128 | 0.5 | 0.5 | 128 | >128 | >128 | >128 | 128 | >128 | 128 | 4 |
| *A. caviae*<br>NUITM-VA2 | 8<br>(1) | 128 | 128 | >128 | 64 | 32 | >128 | >128 | >128 | >128 | >128 | >128 | >128 | 0.25 |
| *E. coli*<br>ATCC 29522 | 0.125<br>(0.125) | 1 | 2 | 2 | 0.25 | 0.032 | 0.125 | 0.5 | 0.008 | 2 | 0.5 | 1 | 0.5 | 0.125 |

Indeed, two copies of *tmexCD3-toprJ1*-like gene clusters and four copies of *tet*(X4) (including the pseudogene) associated with the insertion sequence IS*CR2* were identified in NUITM-VA1, whereas four copies of *tmexCD2-toprJ2*-like gene clusters were identified in NUITM-VA2 (Fig. 1A and B). The identity for *tmexC3, tmexD3*, and *toprJ1* in NUITM-VA1 with the corresponding component genes of *tmexCD3-toprJ1* (accession no. <u>CP066833</u>) (10) were 99.91% (with one amino acid substitution, T235A), 99.90% (with V56E, Q283H, and G591V substitutions), and 100%, respectively. The identity for *tmexC2, tmexD2*, and *toprJ2* in NUITM-VA2 with the corresponding component genes of *tmexCD2-toprJ2* (accession no. <u>CP054471</u>) (9) were 100%, 99.97% (with V56E substitution), and 100%, respectively.

Recently, several subvariants of genes comprised in the *tmexCD-toprJ* cluster were identified, including *tmexD1*.*2* (*tmexD1* variant with V64I substitution) in *Klebsiella pneumoniae* pC5921_mex (IncFIB/IncHI1B/IncU plasmid, accession no. <u>MZ532979</u>) and *tmexD2*.*2* (*tmexD2* variant with V56E and P382A substitutions) in *Klebsiella oxytoca* pC7532_mex (IncFII/IncU plasmid, accession no. <u>MZ532981</u>) (15). In addition, known gene variants of the representative *tmexCD-toprJ* cluster (Table 2), including *tmexC3*.*2* (*tmexC3* variant with Q187H, T256M, and A386T substitutions) and *tmexD3*.*2* (*tmexD3* variant with V610L and L611F substitutions) in *Klebsiella aerogenes* pNUITM-VK5_mdr (IncC/IncX3 plasmid, accession no. <u>LC633285</u>) identified in our previous study (11). Based on these naming-standards, we designated the novel variants of *tmexCD-toprJ* identified in NUITM-VA1 and NUITM-VA2 as *tmexC3*.*3D3*.*3-toprJ1* and *tmexC2D2*.*3-toprJ2*, respectively.

**Table 2.**
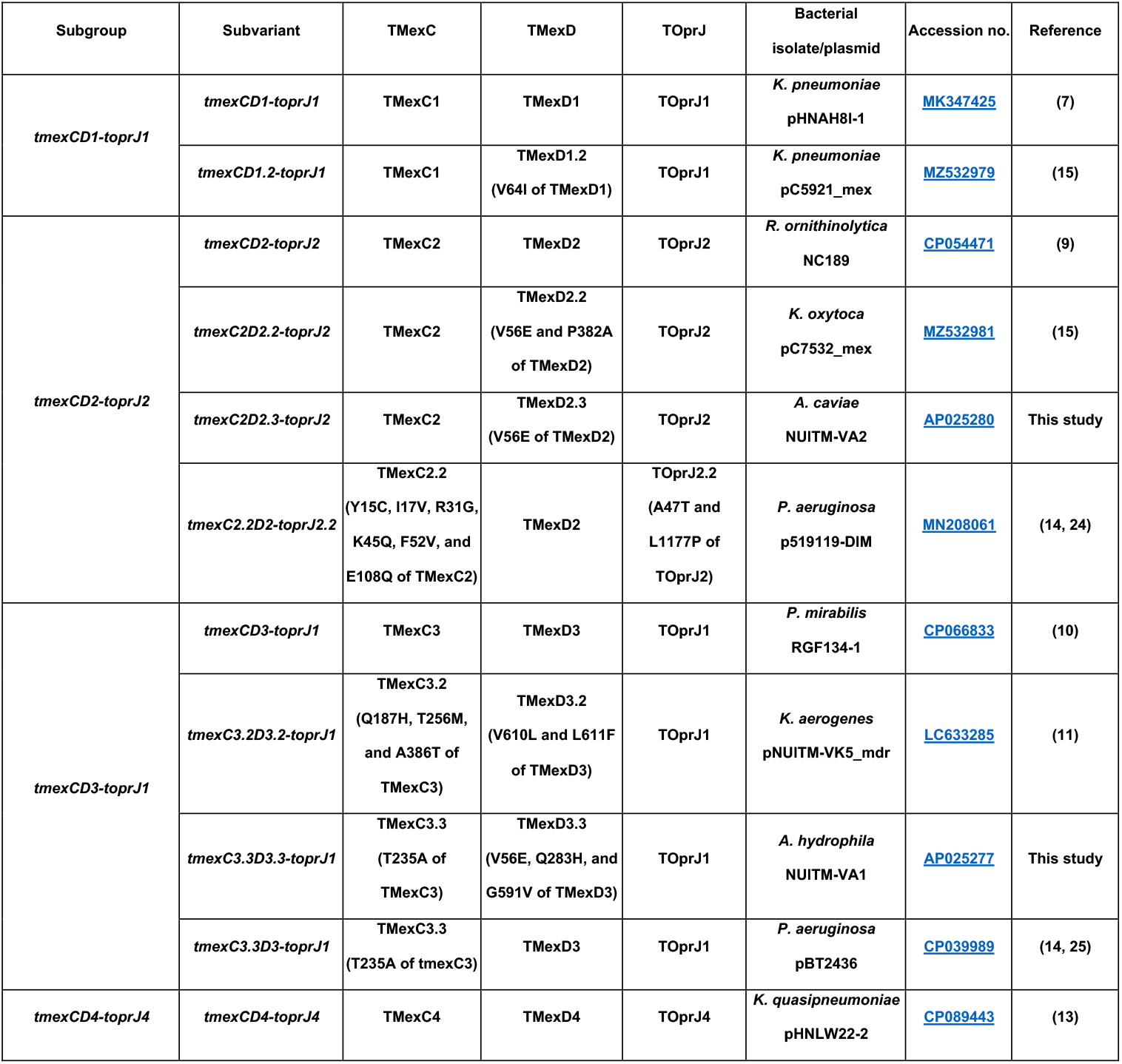
All known variants of mobile RND-type efflux pump gene clusters, *tmexCD-toprJ*. Subgroups and subvariants of *tmexCD-toprJ*, types of component proteins (TMexC, TMexD, and TOprJ), bacterial isolate/plasmid harboring the corresponding *tmexCD-toprJ*, accession nos., and references are shown.

To elucidate the molecular mechanism of intragenomic transposition of *tmexCD-toprJ*, comparative analysis among *tmexCD-toprJ*-containing genomic regions in NUITM-VA1 and NUITM-VA2 was performed. Overall, *tmexCD-toprJ* and the regulator gene *tnfxB* encoded upstream of the gene cluster were found to be associated with an atypical MGE family, SE, that consists of four conserved genes of integrases (*intA* and *intB*), tyrosine recombinase-fold protein (*tfp*), and SE-associated recombination auxiliary protein (*srap*) (17, 18) (Figs. 1C and D).

In NUITM-VA1, one copy of *tmexC3*.*3D3*.*3-toprJ1*-containing SE was flanked by 5′-CATCGA-3′ in *attL* and 5′-TATCGA-3′ in *attR*, whereas the other SE copy was flanked by 5′-TATCGA-3′ and 5′-CATCGA-3′ (Figs. 1A and C). Given the nature of the 6-bp footprint of SE transposition (17, 18), it is likely that one SE copy became the donor for the other SE copy in the second location in NUITM-VA1, but the donor location could not be defined. In NUITM-VA2, four copies of *tmexC2D2*.*3-toprJ2*-containing SEs were detected in the chromosome. The 6-bp fingerprint 5′-TATCGA-3′ was identified next to the 5′-end of the SE copy at the location #1 (4,079,313–4,062,461 nt), as well as next to the 3′-end of the SE copy at the location #2 (3,001,983–3,018,834 nt), and the other SE copy at the location #3 (2,130,980–2,114,127 nt) (Figs. 1B and D). Thus, the SE copy at the location #1 was likely the donor of the other SE copies at locations #2 and #3. Moreover, the 6-bp fingerprint 5′-TATCGA-3′ was identified next to the 5′-end of the SE copy at location #2, as well as next to the 3′-end of the SE copy at the location #4 (1,131,379–1,114,526 nt), suggesting that the SE copy at the location #2 was the donor of the SE copy at the location #4.

Importantly, identical structures as those of *tmexC3*.*3D3*.*3-toprJ1*-containing SE in NUITM-VA1 and *tmexC2D2*.*3-toprJ2*-containing SE in NUITM-VA2 were detected in chromosomes of *A. caviae* WCW1-2 (accession no. <u>CP039832</u>) (19) and *A. caviae* K333 (accession no. <u>CP084031</u>) (20), respectively. Moreover, the *tet*(X4)*-*containing region in NUITM-VA1 was also detected in the WCW1-2 chromosome, suggesting that these genetic structures around ARGs in *Aeromonas* species isolates herein described are not an unusual event.

The known *tmexCD-toprJ* gene cluster can be divided into four major subgroups, consisting of *tmexCD1-toprJ1, tmexCD2-toprJ2, tmexCD3-toprJ1*, and *tmexCD4-toprJ4* (3, 7–16). Hence, the presence of SEs in these *tmexCD-toprJ* subgroups and their subvariants was examined next (Fig. 2). The intact sequences of *tnfxB-tmexCD-toprJ*-containing SEs were identified in all subgroups except *tmexCD4-toprJ4* (Fig. 2A). Although some SEs, such as *tmexCD2-toprJ2*-containing SEs in *Raoultella ornithinolytica* NC189 (accession no. <u>CP054471</u> and <u>MN175502</u>) (9), *tmexC2D2*.*3-toprJ2*-containing SE in *A. caviae* NUITM-VA2 in this study, and *tmexC3D3-toprJ1*-containing SE in *Pseudomonas terrae* subsp. *cibarius* SDQ8C180-2T (accession no. <u>CP073356</u>) (12), harbored insertions of other MGEs within their SEs, such as insertion sequences, the 5’- and 3’-end of the SEs were completely conserved, indicating that they can form functional mobility structures. Interestingly, IncP-2 megaplasmids, such as *P. aeruginosa* pBT2436 (accession no. <u>CP039989</u>) (14, 21), was previously suggested to frequently carry *tmexCD-toprJ*; thus, plasmids also seemed to be involved in such SE-mediated transpositions.

**Fig. 2.**
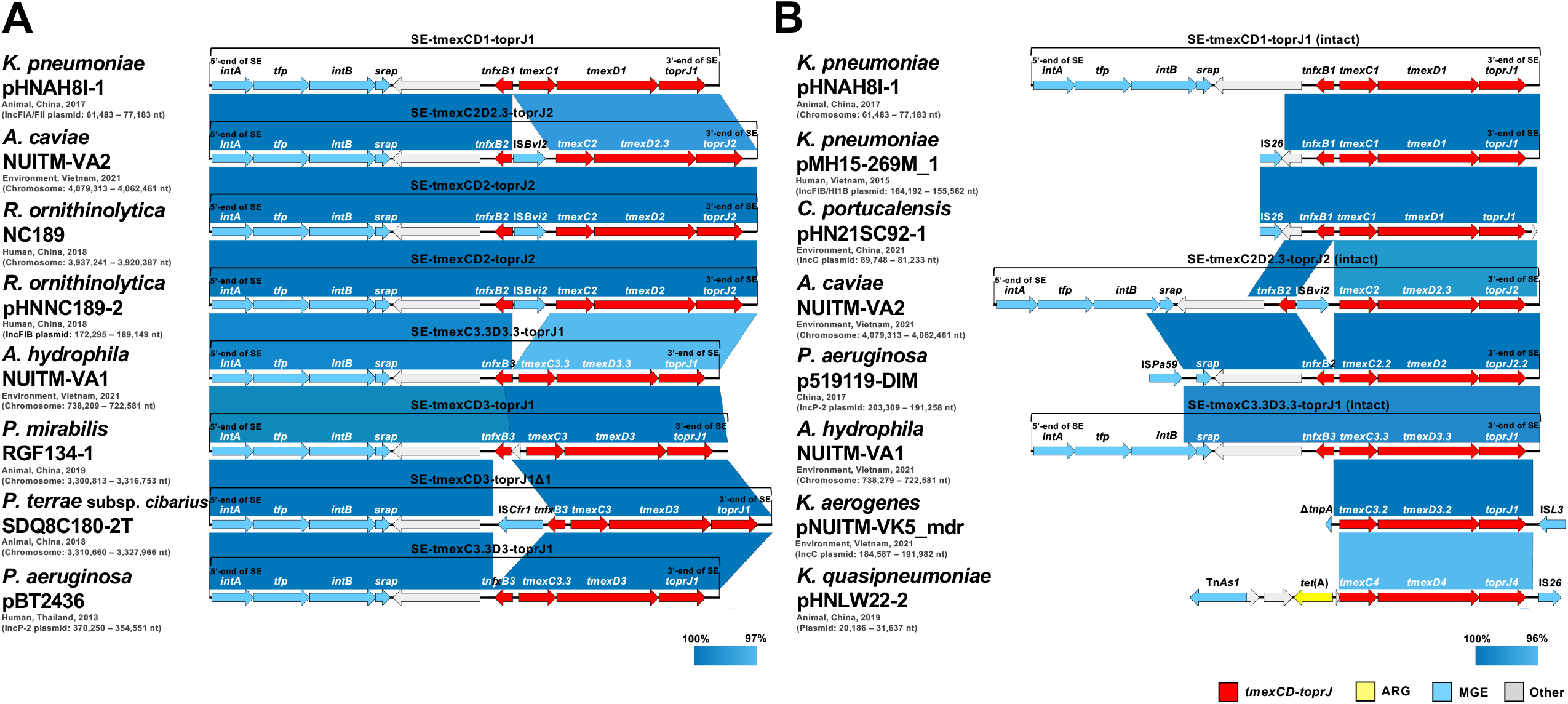
Linear comparison of *tmexCD-toprJ*-containing SEs in *A. hydrophila* NUITM-VA1 and *A. caviae* NUITM-VA2 (accession nos. <u>AP025277</u> and <u>AP025280</u>, respectively) isolated in Vietnam in 2021 in this study with the intact (A) or broken forms (B) in previously reported sequences of *K. pneumoniae* AHM7C8I plasmid pHNAH8I-1 (accession no. <u>MK347425</u>) isolated from an animal in China in 2017 (7), *R. ornithinolytica* NC189 and the plasmid pHNNC189-2 (accession nos. <u>CP054471</u> and <u>MN175502</u>, respectively) isolated from a human in China in 2018 (9), *P. mirabilis* RGF134-1 (accession no. <u>CP066833</u>) isolated from an animal in China in 2019 (10), *P. terrae* subsp. *cibarius* SDQ8C180-2T (accession no. <u>CP073356</u>) isolated from an animal in China in 2018 (12), *P. aeruginosa* T2436 plasmid pBT2436 (accession no. <u>CP039989</u>) isolated from a human in Thailand in 2013 (14, 25), *K. pneumoniae* MH15-269M plasmid pMH15-269M_1 (accession no. <u>AP023338</u>) isolated from a human in Vietnam in 2015 (8), *C. portucalensis* GD21SC92T plasmid pHN21SC92-1 (accession no. <u>CP089438</u>) isolated from an environment in China in 2021 (16), *P. aeruginosa* 1705-19119 plasmid p519119-DIM (accession no. <u>MN208061</u>) isolated in China in 2017 (14, 24), *K. aerogenes* NUITM-VK5 plasmid pNUITM-VK5_mdr (accession no. <u>LC633285</u>) isolated from an environment in Vietnam in 2021 (11), and *K. quasipneumoniae* GLW9C22 plasmid pHNLW22-2 (accession no. <u>CP089443</u>) isolated from an animal in China in 2019 (13). Red, yellow, light blue, and gray indicate *tmexCD-toprJ* gene clusters (*tmexCD-toprJ*), other antimicrobial resistance genes (ARG), mobile gene elements (MGE), and other coding sequences (Other), respectively. The color in comparison of sequences shows the indicated % of identity.

Broken structures close to *tmexCD-toprJ*-containing SEs that lacked the conserved genes of SE and *tmexCD-toprJ*-associated *tnfxB* were also detected (Fig. 2B). This type of broken structures may have concealed the herein described close relationship between *tmexCD-toprJ* and SEs in previous studies. For example, *tmexCD1-toprJ1* in *K. pneumoniae* pMH15-269M_1 (IncFIB/HI1B plasmid, accession no. <u>AP023338</u>) (8) and *Citrobacter portucalensis* pHN21SC92-1 (IncC plasmid, accession no. <u>CP089438</u>) (16) was previously suspected of being mobilized by IS*26*, but intact 3’-end sequence of SE was present on pMH15-269M_1 but not on pHN21SC92-1, suggesting the involvement of other MGEs rather than SE is an event that occurred after initial SE-mediated transpositions. For *tmexCD4-toprJ4* in *Klebsiella quasipneumoniae* pHNLW22-2 (untypeable plasmid, accession no. <u>CP089443</u>) (13), no evidence suggested SE involvement on either the 3’ or 5’ sides of the gene cluster; nevertheless, since this *tmexCD-toprJ* variant has only been reported in one case to date, a possible association with SE cannot be ruled out.

In conclusion, our present study provides a glimpse into *Aeromonas* species, one of the most common environmental bacteria that have been rapidly and silently becoming resistant to clinically important antimicrobials. Notably, our present study highlights for the first time the role of SE-mediated transpositions for the evolution of the MDR gene clusters, *tmexCD-toprJ*. Our previous study suggested that *Aeromonas* and *Pseudomonas* species in the natural environment are not only important reservoirs of ARGs, but are also carriers of evolutionary changes in ARGs, such as *bla*_GES-5_-like carbapenemase genes (22). The MGE-mediated spread of ARGs among bacteria and their epidemiology concerning some specific examples, such as *mcr1* and IS*Apl1* (23), and *bla*_NDM_ and Tn*125* (24), have been previously analyzed in detail. The present study provides more direct epidemiological evidence of transpositions of *tmexCD-toprJ* mediated by SEs, which can be identified by the footprints of SEs. This finding is quite significant for investigating the global spreading of ARGs, including *tmexCD-toprJ*, and will pave the way for future genomic epidemiological investigations on antimicrobial-resistant bacteria.

## Materials and methods

### Bacterial isolation and antimicrobial susceptibility testing

Carbapenem- and tigecycline-resistant of environmental isolates of *Aeromonas* species, *A. hydrophila* NUITM-VA1 and *A. caviae* NUITM-VA2 were obtained from the Kim-Nguu River in Hanoi, Vietnam, in March 2021. Environmental water sample was collected and cultured using Luria-Bertani (LB) broth containing 4 mg/L of meropenem at 37°C overnight, and then further selected and isolated using CHROMagar COL-APSE (CHROMagar Microbiology) containing 4 mg/L of tigecycline. Bacterial species identification was performed using MALDI Biotyper (Bruker). Antimicrobial susceptibility testing using *Escherichia coli* ATCC 25922 as quality control was performed with agar dilution according to the CLSI 2020 guidelines. For tigecycline, AST was additionally performed in the presence or absence of 75 mg/L of the efflux pump inhibitor 1-(1-naphthylmethyl)-piperazine (NMP) as used in the previous studies (7, 11).

### Whole-genome sequencing and subsequent bioinformatics analysis

Whole-genome sequencing of NUITM-VA1 and NUITM-VA2 was performed using MiSeq (Illumina) with MiSeq Reagent Kit v2 (300-cycle) and MinION (Oxford Nanopore Technologies; ONT) with the R9.4.1 flow cell. The library for Illumina sequencing (paired-end, insert size of 300–800 bp) was prepared using the Nextera XT DNA Library Prep Kit, and the library for MinION sequencing was prepared using the Rapid Barcoding Kit (SQK-RBK004). ONT reads were basecalled using Guppy v5.0.11 in the super-accuracy mode and then assembled de novo using Canu v2.1.1 (https://github.com/marbl/canu) with the default parameters. The overlap regions in the assembled contigs were detected using LAST (https://gitlab.com/mcfrith/last) and then trimmed manually. Sequencing errors were corrected by Racon v1.4.20 (https://github.com/isovic/racon) twice with the default parameters using ONT reads and then corrected by Pilon v1.20.1 (https://github.com/broadinstitute/pilon) twice with the default parameters using Illumina reads, resulting in their complete circular chromosomes.

Genome annotation and average nucleotide identity analyses were performed using the DFAST server (https://dfast.nig.ac.jp). ARGs were detected using Staramr v0.7.2 (https://github.com/phac-nml/staramr) with the custom ARGs database, including all known tigecycline resistance genes. The circular representation of bacterial chromosomes was visualized using the Proksee server (https://proksee.ca). Linear comparison of sequence alignments of genomic regions containing ARGs and MGEs was performed using BLAST and visualized by Easyfig v2.2.2 (http://mjsull.github.io/Easyfig/).

### Nucleotide sequence accession nos

Complete chromosome sequences of *A. hydrophila* NUITM-VA1 and *A. caviae* NUITM-VA2 have been deposited at GenBank/EMBL/DDBJ under accession nos. <u>AP025277</u> and <u>AP025280</u>, respectively.

## Ethical approval

Not required.

